# Genomic inferences of domestication events are corroborated by written records in *Brassica rapa*

**DOI:** 10.1101/118372

**Authors:** Xinshuai Qi, Hong An, Aaron Ragsdale, Tara E. Hall, Ryan N. Gutenkunst, J. Chris Pires, Michael S. Barker

**Affiliations:** Department of Ecology & Evolutionary Biology, University of Arizona, Tucson, Arizona, USA 85721; Division of Biological Sciences, University of Missouri, Columbia, Missouri, USA 65211; National Key Lab of Crop Genetic Improvement, Huazhong Agricultural University, Wuhan, P. R. China 430070; Program in Applied Mathematics, University of Arizona, Tucson, Arizona, USA 85721; Department of Molecular and Cellular Biology, University of Arizona, Tucson, Arizona, USA 85721

**Keywords:** *Brassica rapa*, genetic structure, transcriptome, domestication, crop improvement, population genomics

## Abstract

Demographic modeling is often used with population genomic data to infer the relationships and ages among populations. However, relatively few analyses are able to validate these inferences with independent data. Here, we leverage written records that describe distinct *Brassica rapa* crops to corroborate demographic models of domestication. *Brassica rapa* crops are renowned for their outstanding morphological diversity, but the relationships and order of domestication remains unclear. We generated genome-wide SNPs from 126 accessions collected globally using high-throughput transcriptome data. Analyses of more than 31,000 SNPs across the *B. rapa* genome revealed evidence for five distinct genetic groups and supported a European-Central Asian origin of *B. rapa* crops. Our results supported the traditionally recognized South Asian and East Asian *B. rapa* groups with evidence that pak choi, Chinese cabbage, and yellow sarson are likely monophyletic groups. In contrast, the oil-type *B. rapa* subsp. *oleifera* and brown sarson were polyphyletic. We also found no evidence to support the contention that rapini is the wild type or the earliest domesticated subspecies of *B. rapa.* Demographic analyses suggested that *B. rapa* was introduced to Asia 2400-4100 years ago, and that Chinese cabbage originated 1200-2100 years ago via admixture of pak choi and European-Central Asian *B. rapa.* We also inferred significantly different levels of founder effect among the *B. rapa* subspecies. Written records from antiquity that document these crops are consistent with these inferences. The concordance between our age estimates of domestication events with historical records provides unique support for our demographic inferences.

## Introduction

Demographic analyses are widely used to infer the history of populations. Many analyses make demographic inferences of ancient events that occurred in prehistory. Thus, there is often no opportunity to validate these inferences with independent data. Domestication offers a unique chance to verify the inferences from methods that are used in model and non-model species. Artificial selection and domestication have long served as a testing ground for understanding evolution (Darwin 1859; Diamond 2002). The study of crop domestication provides fundamental insights on crop history and improvement, but also contributes some of our most well understood systems to reveal the powers and limits of selection to shape organisms (Ross-Ibarra *et al.* 2007; Purugganan & Fuller 2009; Gross & Olsen 2010; Olsen & Wendel 2013; Meyer & Purugganan 2013). *Brassica* crops are arguably one of the best examples to illustrate the diversity that can be created by artificial selection (Purugganan *et al.* 2000; Prakash *et al.* 2011). The three *Brassica* crops most well known for their morphological diversity are *B. rapa* (AA, 2n = 20), *B. oleracea* (CC, 2n = 18), and their allopolyploid derivative, *B. napus* (AACC, 2n = 38). Each of these taxa includes several cultivated subspecies that were domesticated for a variety of uses. Long considered a classic textbook example of plant domestication (Darwin 1859; Ladizinsky 2012; Zohary *et al.* 2012) and the power of artificial selection (Purugganan *et al.* 2000), *B. oleracea* includes a diversity of crops such as broccoli, cauliflower, cabbage, kale, Brussels sprout, and kohlrabi. However, its sister species, *B. rapa,* has similar morphological variation and represents a parallel example for comparative analyses of artificial selection and evolution. Several subspecies are recognized in *B. rapa* that have been cultivated for particular phenotypic characteristics. These include turnip (subsp. *rapa*) with an enlarged edible root, yellow (subsp. *trilocularis*) and brown sarsons (subsp. *dichotoma*) that are used as mustard and oil seeds mainly in India, the oilseed field mustard (subsp. *oleifera*), and the leafy East Asian vegetables pak choi (subsp. *chinensis*) and Chinese cabbage (subsp. *pekinensis*) with their enlarged mid-rib and leaves (Fig. 1). There are also several less broadly known leafy subspecies, including rapini (subsp. *sylvestris),* choy sum (subsp. *parachinensis*), zi cai tai (subsp. *purpuraria*), tatsoi (subsp. *narinosa*), mizuna (subsp. *nipposinica*), and komatsuna (or Japanese mustard spinach/rapini, subsp. *perviridis*).

**Fig. 1.**
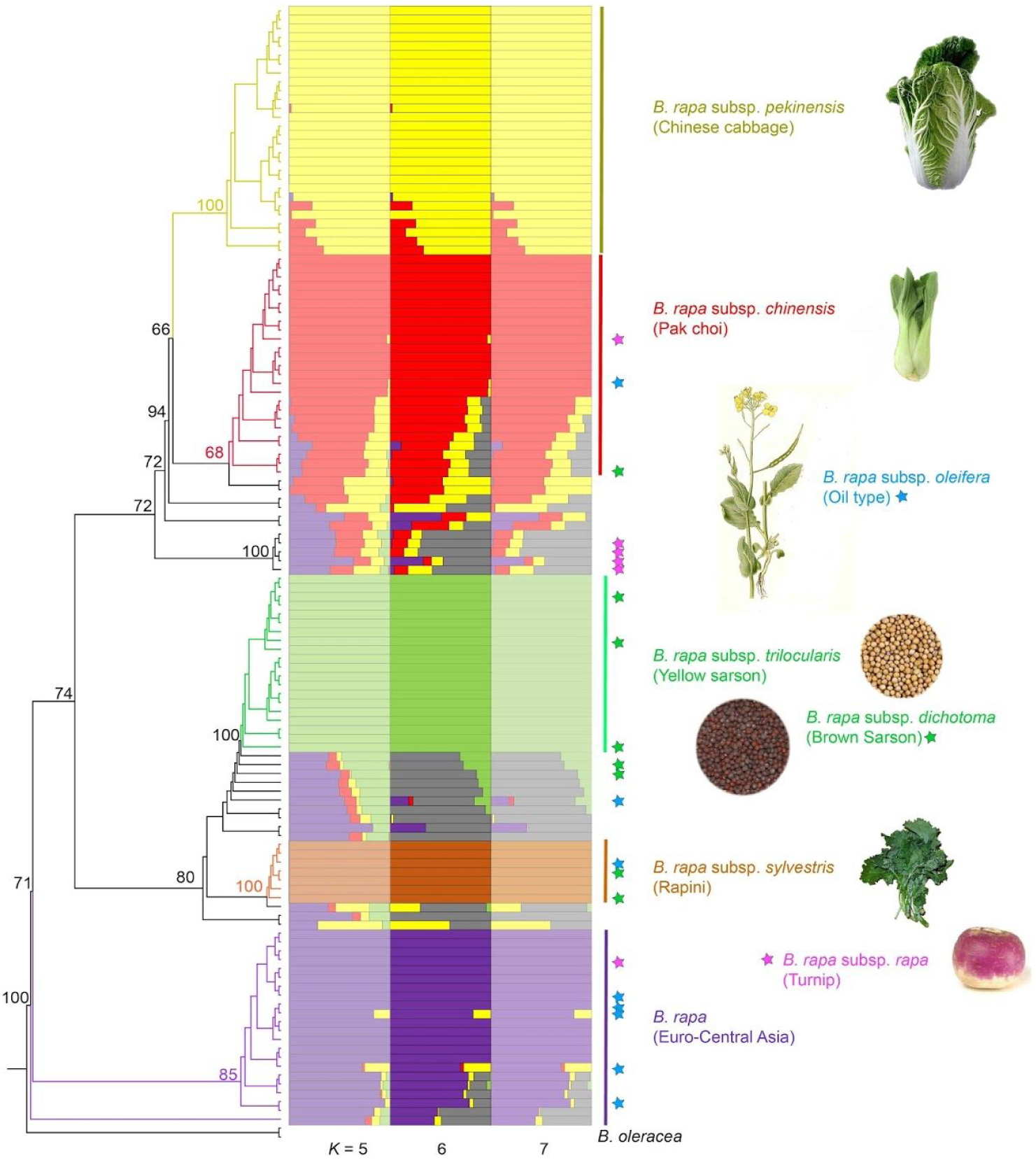
The integrated maximum likelihood phylogeny and fastStructure results of the 126 *Brassica rapa* accessions analyzed here. The five *B. rapa* genetic groups are represented by Europe-Central Asian *B. rapa,* rapini (subsp. *sylvestris*), yellow sarson (subsp. *trilocularis*), pak choi (subsp. *chinensis*), and Chinese cabbage (subsp. *pekinensis*). fastStructure ancestry proportions shown from *K* = 5 to 7 with optimal *K* = 6. Blue, green, and pink stars represent *B. rapa* subsp. *oleifera,* subsp. *dichotoma,* and subsp. *rapa* accessions. Two *B. oleracea* accessions were used to root the phylogenetic tree. Bootstrap support values of key nodes are labeled on the tree. Attributions for *B. rapa* images are in Table S4.

Domestication of the diverse *B. rapa* crops most likely occurred during written history (after ~3500 BCE). Many of the classically studied crops, such as maize and Asian rice, were domesticated in prehistory (Smith 1997; Piperno & Flannery 2001; Long *et al.* 2006; Kovach *et al.* 2007; Piperno *et al.* 2009; Fuller *et al.* 2009; Molina *et al.* 2011; Sang & Ge 2013; Gross & Zhao 2014). A written record of domestication for most crops is simply unavailable to develop hypotheses and corroborate genomic analyses. In contrast, the oldest mention of *Brassica* crops in general dates back nearly 5000 years. *Brassica* crops were first described in a Chinese almanac from ~3000 BCE and ancient Indian texts from around 1500 BCE (Prakash *et al.* 2011). Although the record is unclear on which European *B. rapa* crop was first domesticated, turnips were on the list of plants grown in the garden of Merodach-baladan in Babylonia (722-711 BCE) (Körber-Grohne 1987). Consistent with this early written record, archaeological evidence of turnip seeds and roots have been reported from multiple archeological sites of Neolithic villages in Switzerland, India, and China (Prakash and Hinata 1980; Ignatov et al. 2008). *Brassica rapa* subsp. *rapa* was clearly described nearly 2600 years ago in Shi Jing, the oldest existing Chinese classic poetry collection (Li 1981; Ye 1989; Luo 1992). The authors of Shi Jing used the ancient Chinese character for turnip, indicating that this crop was in East Asia by that time. The earliest historical records of Chinese cabbage (*B. rapa* subs, *pekinensis*), “Niu Du Song (beef tripe cabbage)”, appears in Xin Xiu Ben Cao, one of the earliest official pharmacy books edited by Jing Su (659 CE). A clear description of Chinese cabbage is also found in Ben Cao Tu Jing (1061 CE) by Song Su. He described Niu Du Song in Yangzhou city in Southern China as a plant with large, round, and tender leaves with a uniquely wrinkled leaf. Based on these documents, we may expect that many of the the Asian *B. rapa* crops were domesticated as long as 3500 years ago and that *B. rapa* subsp. *pekinensis* may be at least 1500 years old.

Despite the economic importance and historical records of *B. rapa* crops, our understanding of their relationships and origins is still not well resolved. Previous studies have attempted to identify genetically discrete groups among *B. rapa* crops with molecular markers (McGrath & Quiros 1992; Ren *et al.* 1995; Zhao *et al.* 2005, 2007, 2010; Takuno *et al.* 2006; Del Carpio *et al.* 2011a; b; Guo *et al.* 2014; Pang *et al.* 2015; Takahashi *et al.* 2015; Tanhuanpää *et al.* 2016). Most analyses identify three ambiguous geographic groups present in Europe and Central Asia, South Asia, and East Asia (Del Carpio *et al.* 2011a; b; Guo *et al.* 2014; Pang *et al.* 2015) except a recent analysis that found no evidence for geographic structure among diverse *B. rapa* accessions (Tanhuanpää *et al.* 2016). In addition, previous genetic and empirical studies suggested two distinct hypotheses for the origin of Chinese cabbage: domesticated directly from pak choi (Song *et al.* 1990; Zhao *et al.* 2005; Takuno *et al.* 2006) or via genetic admixture of turnip and pak choi(Li 1981; Ren *et al.* 1995). Although many contemporary studies have included large and diverse samples, the major limitation to distinguishing *B. rapa* subspecies and crops may be the number and quality of the genetic markers. Fine scale resolution of relationships among groups within *B. rapa* may be beyond the power of traditional marker systems because of their relatively short evolutionary history with high rates of outcrossing. Clarification of the number of genetically distinct groups and their relationships is needed to address basic questions about the domestication of *B. rapa.*

Genome assemblies of multiple *Brassica* species (Wang *et al.* 2011; Liu *et al.* 2014; Parkin *et al.* 2014; Chalhoub *et al.* 2014) and advancements in sequencing technology provide an opportunity to resolve the relationships among subspecies and the domestication history of *B. rapa.* Recently, Cheng et al. (2016) resequenced 199 *B. rapa* accessions. Their STRUCTURE and phylogenomic analyses provided improved support for the relationships among the primary subspecies within *B. rapa.* However, their sampling did not include several important Eurasian *B. rapa* crops that are crucial for understanding the timing and order of *B. rapa* domestication. For example, no samples were collected from Central Asia, the potential center of origin for *B. rapa.* Cheng et al. (2016) also only sampled one yellow sarson accession and did not include brown sarson (subsp. *dichotoma)* or rapini (subsp. *sylvestris).* Diverse sampling of accessions combined with population genomic approaches are needed to better resolve the complex series of events that occurred during domestication of the *B. rapa* crops.

In this study, we sequenced the transcriptomes of 126 accessions representative of the geographic and crop diversity of *B. rapa.* We analyzed thousands of SNPs from across the nuclear genome to resolve the genetic structure and relationships of *B. rapa* crops. Using a diffusion approximation approach, ∂ a ∂ i (Gutenkunst *et al.* 2009), we evaluated models of the demographic history and diversification of *B. rapa,* as well as estimated the timing of domestication events in China and India. These age estimates were compared to the written records available for each phase of *B. rapa* domestication. We also estimated the reduction of genetic diversity and effective population size in each Asian *B. rapa* subspecies due to the potential founder effect during their introduction and domestication in Asia. Our analyses provide new insights into the origins and relationships of *B. rapa* crops. Given that most studies of domestication are focused on crops that arose during prehistory, our analyses of *B. rapa* provide a unique opportunity to explore crop domestication and diversification with a written record.

## Materials and Methods

### Plant Materials

We collected seeds of 143 *B. rapa* accessions, with 139 samples from USDA GRIN database (http://www.ars-arin.aov/1 and four from the authors’ collection. Samples were selected to maximize representation of major *B. rapa* subspecies. These collections include *B. rapa* and its 11 subspecies from 20 countries worldwide (Table S1). Six of the major subspecies have more than eight representatives, randomly chosen from different sources, with the exception of the two *sylvestris* lines collected by Dr. Gómez Campo. The Asian vegetable subspecies *parachinensis, narinosa, nipposinica* and *perviridis* are likely derived from either or both *chinensis* and *pekinensis* (Cheng et al. 2016). Therefore, only representative accessions of these Asian vegetable subspecies were included in this analysis. After we removed misidentified seed accessions (see below), 126 *B. rapa* accessions remained for the following analyses.

### Plant Growth Conditions and Tissue Collection

All seeds were grown in a Conviron growth chamber with 16 hours of light at 23°C. and 8 hours of dark at 20°C. at the Bond Life Science Center, University of Missouri (Columbia, Missouri, USA) during March to May 2015. All samples were taken from the second youngest leaf 20 days after germination, then snap-frozen in liquid nitrogen and stored at -80°C.

### Sample Preparation, RNA Isolation, Library Construction

Total RNA was isolated using the PureLinkTM RNA Mini Kit (Invitrogen, USA), following the instructions. RNA samples were qualified and quantified by a Nanodrop 1000 spectrophotometer (Nanodrop Technologies, USA). After that, double-stranded cDNA were synthesized according to the manufacturer’s protocol of the Maxima H Minus Double-Stranded cDNA Synthesis Kit (Thermo, Lithuania) and purified by the GeneJET PCR Purification Kit (Thermo, Lithuania). All samples were sequenced with Illumina HiSeq 2000 system at the University of Missouri DNA core. Each sample contained about 9 million 250 bp pair-end reads. SRA files for the samples are available at NCBI-SRA (SRP072186, http://www.ncbi.nlm.nih.gov/sra/SRP0721861.

### Sequence Assembly and Alignment

We first trimmed each fastq file using Trimmomatic (Bolger *et al.* 2014). Read quality was assessed with FASTQC (Andrews & Others 2010). The reference *B. rapa* genome (version 1.5) was downloaded from the Brassica Database (http://brassicadb.ora/brad/1).

For the reference based assemblies, we used Tophat version 2.0.14 (Trapnell *et al.* 2009) with Bowtie2 version 2.2.5 (Langmead & Salzberg 2012). Variant calling was performed using Samtools (Li *et al.* 2009). We retained only SNPs and did not consider indels or MNPs in subsequent analyses. To obtain high quality SNPs for downstream analyses, we performed multiple SNP filtering steps: the SNPs were first filtered with *vcfutils.pl* script in SAMtools/bcftools package (Li *et al.* 2009) and *vcffilter* to only include SNPs with depth greater than 10 and mapping quality greater than 30. For the phylogenetic and genetic structure analyses, we then filtered the remaining SNPs with PLINK version 1.9 (Purcell *et al.* 2007) to only include SNPs with a > 90% genotyping rate and minor allele frequency (MAF) higher than 5%. In contrast, our filtering for the ∂ a ∂ i demographic models and nucleotide diversity analyses retained many more SNPs to estimate the allele frequency spectrum. Following filtering for depth and mapping quality, we annotated the SNPs with SnpEff (Cingolani *et al.* 2012) and only retained synonymous SNPs. Further, we implemented a 25 kb sliding window neutrality test (Tajima’s D) in vcftools (Danecek *et al.* 2011) to filter out SNPs that may be directly or indirectly under selection.

### Identification of Misidentified *Brassica* Seed Contaminants

Misidentified seed accessions are not uncommon in plant germplasm collections and can confound analyses if not recognized. To identify potential seed contamination from *B. napus* and *B. oleracea,* we BLASTed the *de novo* assembled reading frames of each newly sequenced accession against the latest *B. rapa, B. oleracea* (TO1000 version 2.1; genomevolution.org) and *B. napus* (version 4.1; brassicadb.org) genomes. *De novo* assemblies for each accession were performed in SOAPdenovo-Trans version 1.03 (Xie *et al.* 2014) with a 127 kmer. Accessions with a majority of best BLAST hits to *B. oleracea* and/or *B. napus* were flagged as potential contaminants. We further verified the suspicious accessions through DNA C-value measurement using flow cytometry. Fresh leaf samples were sent to the Flow Cytometry Lab at the Benaroya Research Institute (WA) for flow cytometry. The C-values of *B. rapa, B. oleraceae,* and *B. napus* are significantly different from each other and provide an alternative method of verification. A total of 11 suspicious *B. napus,* two suspicious *B. oleracea* and four suspicious autopolyploid *B. rapa* were identified among the *B. rapa* accessions from the USDA germplasm seed bank (Table S2). These accessions were removed from further analyses in this study. After contaminant removal, 126 *B. rapa* accessions remained for the following analyses.

### Phylogenetic Inference and Genetic Structure

SNPs called from mapping reads to the *Brassica rapa* reference genome were used for phylogenetic inference and genetic structure analyses. The SNP dataset was concatenated from ped format to fasta format using a custom perl script. The data set included SNPs from our 126 *B. rapa* accessions as well as two *B. oleracea* as outgroups (SRA accession: SRR1032050 and SRR630924, http://www.ncbi.nlm.nih.gov/sra). A maximum likelihood phylogeny was constructed using RAxML version 8.2.4 (Stamatakis 2014). We implemented RAxML’s rapid bootstrap algorithm with the GTR+GAMMA model and rooted the phylogeny using the two *B. oleracea* accessions. Genetic structure among the accessions was inferred using a Bayesian clustering method implemented in fastStrucutre (Raj *et al.* 2014). We tested *K* = 1 to 10 clusters with five replicates for each *K* using the default convergence criterion and prior. The optimal *K* value was estimated with the *chooseK* tool contained in the fastStructure package. The results of all replicates were summarized using CLUMPAK (http://clumpak.tau.ac.il/)(Kopelman *et al.* 2015). The multiple *K* value fastStructure results were re-ordered and visualized in Excel based on the sample order of the phylogenetic inference.

### Inference of Demographic History with ∂ a ∂ i

Models of the demographic history of the major *B. rapa* groups were evaluated using diffusion approximations to the allele frequency spectrum (AFS) with ∂ a ∂ i (Gutenkunst *et al.* 2009). This approach allowed us to rapidly and flexibly test alternative demographic models.

Only individuals that could be readily attributed to one of the major genetic lineages identified in our fastStructure and phylogenetic analyses were used in this analysis (denoted by the vertical bars on Figure 1). Individuals were assigned a group if they had >50% fastStructure identity and were phylogenetically clustered. We analyzed the demographic history of 102 *B. rapa* accessions divided into five groups (Table S1): 22 accessions of European and Central Asian *B. rapa* (group “EU-CA”), 7 accessions representing *B. rapa* subsp. *sylvestris* (group “S”), 20 accessions of *B. rapa* subsp. *trilocularis* and *B. rapa* subsp. *dichotoma* (group “**I**”), 25 accessions representing *B. rapa* subsp. *chinensis* (group “C”), and 28 accessions of *B. rapa* subsp. *pekinensis* (group “P”). Based on the phylogeny, we also combined groups S and I as a South Asian group “SA” in some analyses. Likewise, groups C and P were merged in some analyses as an East Asian group “EΑ”.

We implemented a model testing hierarchy to accommodate ∂ a ∂ i’s limit of analyzing three populations at a time. We first evaluated one- and two-population models for each of the five *B. rapa* groups to estimate the timing of divergence among the derived groups. The results of these initial analyses informed our subsequent demographic modeling of three different three-population model groups (Fig. 2; Table 1) described in further detail below. Notably, we also performed the same simulations under exponential growth models, but the models did not converge (results not shown).

**Table 1.**
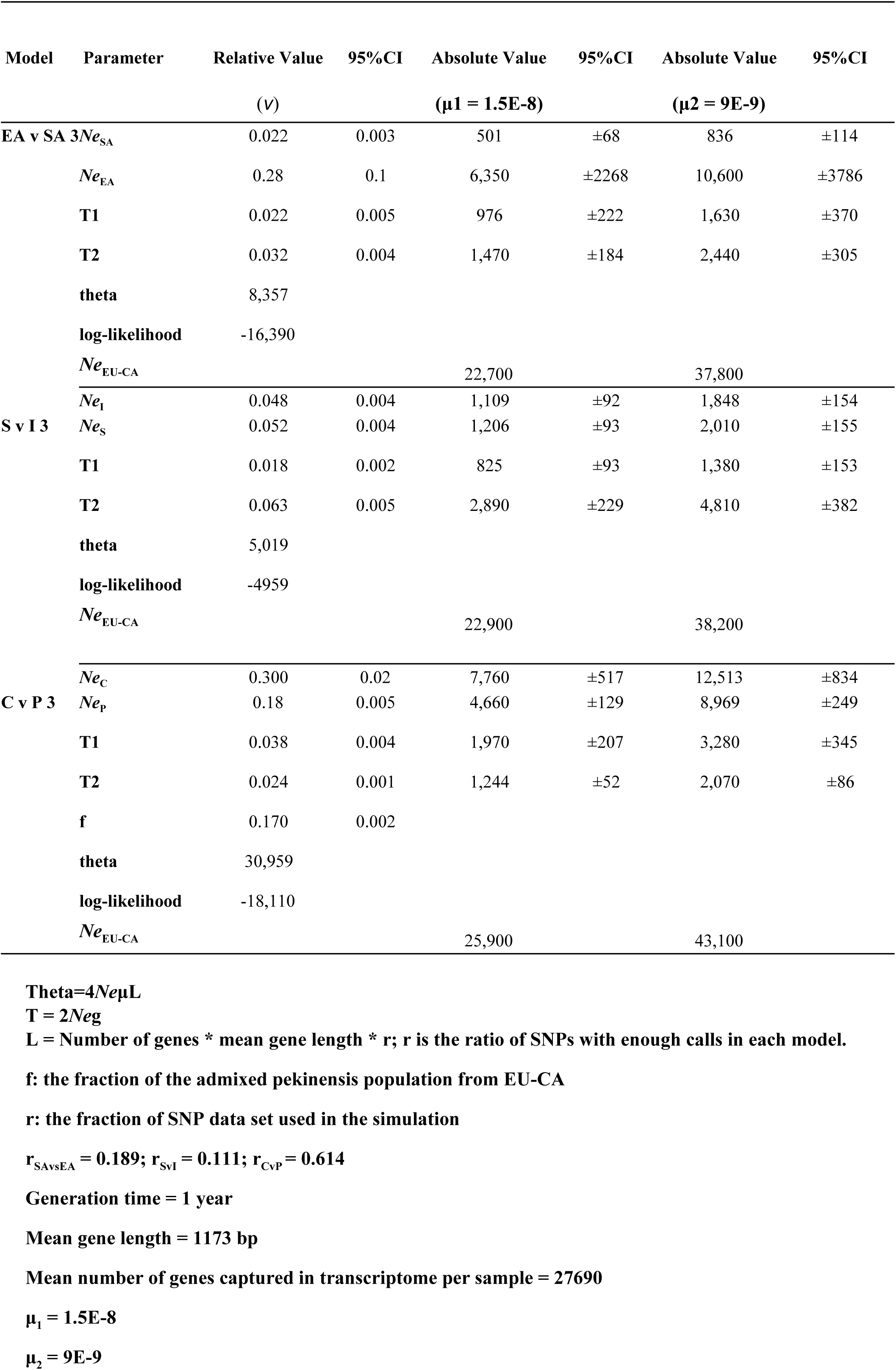
The parameters and confidence intervals inferred in ∂ a ∂ i simulations. These parameters correspond to those displayed in Fig 3. The 95% Cl was calculated using the Godambe bootstrapping method. Unit of absolute effective population size: individual; unit of absolute time: year.

**Fig. 2.**
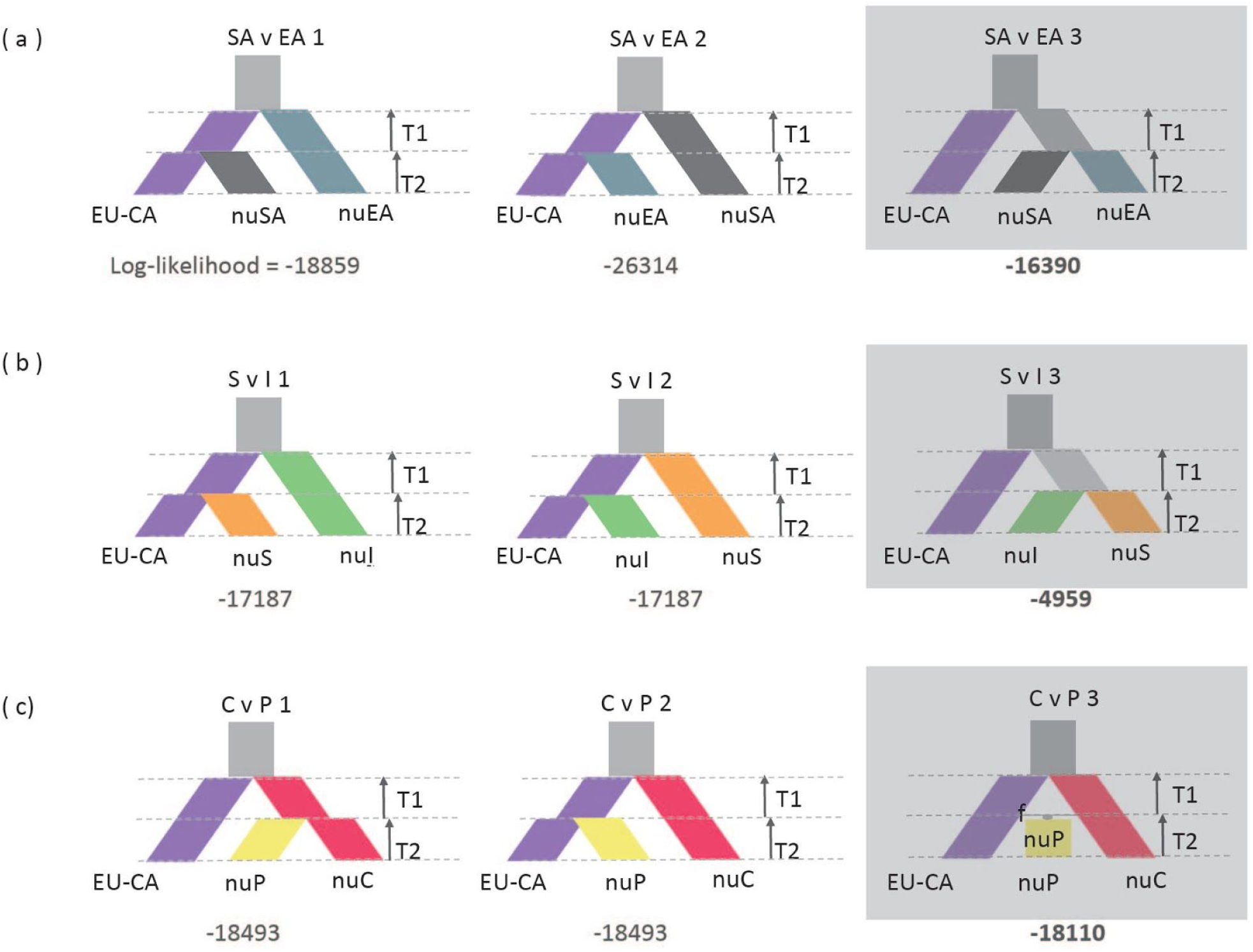
The demographic models evaluated with *∂* a ∂ i. The models with gray background color are the optimal model under each scenario. The Log-likelihood value is indicated under each model. EU-CA represents *B. rapa* and *B. rapa* subsp. *rapa* group; SA represents *B. rapa* subsp. *sylvestris, B. rapa* subsp. *trilocularis* and *B. rapa* subsp. *dichotoma* group; EA represents *B. rapa* subsp. *chinensis* and *B. rapa* subsp. *pekinensis* group; S represents *B. rapa* subsp. *sylvestris* group; I represents *B. rapa* subsp. *trilocularis* and *B. rapa* subsp. *dichotoma* group; C represents *B. rapa* subsp. *chinensis* group; P represents *B. rapa* subsp. *pekinensis* group.

#### Model Group A

In model group A, we tested whether the SA and EA groups were independently derived from EU-CA, or whether a group ancestral to both SA and EA split from EU-CA and then split into SA and EA (Fig. 2a).

#### Model Group B

Model group B focused on the origins of S and I. We tested whether S and I independently split from EU-CA, or if a common ancestor of S and I split from EU-CA and then subsequently diverged into S and I (Fig. 2b).

#### Model Group C

The models in group C evaluated long debated alternative hypotheses on the origin of Chinese cabbage, *B. rapa* subsp. *pekinensis* (Fig. 2c). Especially as to whether it was originated from *chinensis* (Song *et al.* 1990; Zhao *et al.* 2005; Takuno *et al.* 2006), from EU-CA rapa, or from the admixture of the first two (Li 1981; Ren *et al.* 1995).

In each model, the EU-CA population was assumed to have constant effective population size. Each of these models was parameterized by the timing between each split and admixture event (e.g. T1, T2) and the relative effective population sizes over those time intervals (e.g. vEA, vSA). The effective population sizes were relative to the ancestral population size, so *v* less than one indicates a population size decline, while *v* greater than one indicates growth. Under the third *pekinensis* origin model (C v P 3; Fig. 2C), an additional parameter f (between 0 and 1) estimated the fraction of the admixed *pekinensis* population from EU-CA.

#### Model Optimization and Testing

Our filtered SNP collections were converted to the ∂ a ∂ i frequency spectrum file format using a publicly available perl script (https://github.com/owensgl/reformat/blob/master/vcf2dadi.pl). In order to correct for linkage among the SNPs, we performed a likelihood ratio test on the models for each population using the Godambe Information Matrix (GIM)(Godambe, 1960). This test statistic is compared to a χ^2^ distribution with degrees of freedom equal to the difference in number of parameters between the simple and complex model. The linkage of SNPs requires us to adjust the test statistic, which is calculated via the GIM. We constructed the three-population folded joint frequency spectra by projecting down to a specified sample size for each population (Marth *et al.* 2004). Because many SNPs were not called in every sequenced individual, by projecting the frequency spectrum to a smaller sample size, we were able to include more SNPs in our analysis. The sample size that we projected to in each population was chosen so that the frequency spectrum maximized the total number of segregating sites for each population. We projected to 20, 40, and 40 samples for the EU-CA, SA, and EA populations in the model group A analysis, respectively. For model group B, we projected to 13, 40, and 14 samples for the EU-CA, *trilocularis,* and *sylvestris* populations, resp. For model group C, we projected to 11, 13, and 23 for the EU-CA, *chinensis,* and *pekinensis* populations. For *B. rapa* subsp. *sylvestris, trilocularis,* and *dichotoma,* many SNPs for which genotypes were missing from some individuals showed a large excess of heterozygotes, and this pattern was not seen for SNPs called in every individual. Thus, we used SNPs called in every individual in the *sylvestris, trilocularis,* and *dichotoma* group to avoid any artifacts in the projected frequency spectrum.

During optimization we imposed lower and upper bounds on model parameters, with times allowed between 0.001 and 1 genetic time units, and population size changes between 0.001 and 100 of the reference (ancestral) population size. We used built in maximum likelihood optimization in ∂ a ∂ i to fit the expected frequency spectra under the parameterized models to the observed spectra. The population scaled mutation rate θ was a free parameter of the models. We repeated the optimization for each model 100 times from random initial parameter guesses, to ensure consistent convergence to the optimal parameters. The Godambe bootstrapping method (Coffman *et al.* 2016) was performed to estimate the 95% confidence intervals of the best fit parameters. Bootstrap datasets were generated with a custom python script.

We selected the best fitting model for each group of models by selecting the model with highest log-likelihood under the best-fit parameter set. For the best fitting models in each group, we converted the inferred population genetic parameters to years using the relation θ = *4NeμL* and T = 2Neg. L is the effective length in base pairs of the region from which the SNP data was obtained. L was calculated as the product of mean gene length, total genes covered by the SNP dataset, and the fraction of SNP data set used in the simulation due to missing data. Upper and lower mutation rates of *μ* = 1.5 **×** 10^−8^ and 9 **×** 10^−9^ per synonymous site per generation and generation time of one year were used in our age estimates (Koch *et al.* 2000; Kagale *et al.* 2014).

### Genetic Diversity Analysis

To assess the level of genetic diversity of the five identified genetic groups, we estimated the average nucleotide diversity, π (Nei & Li 1979), using vcftools version 0.1.12 (Danecek *et al.* 2011). A window size of 100 kbp and step size of 25 kbp were applied to the genetic diversity estimation. All calculations and visualizations were made in R (version 3.2.2) and Rstudio (Version 0.99.482).

## Results

### Read Mapping and SNP Filtering

Using paired-end Illumina RNA-seq, we sequenced the transcriptomes of 143 *Brassica rapa* accessions with 250 bp mean read length. Following the removal of 17 misidentified accessions were not clearly *B. rapa* (Table S2), the transcriptomes of 126 *B. rapa* accessions were used in our analyses. On average, 8.09 million reads per sample were left after Trimmomatic read cleaning. For each of the reference genome based assemblies, 33.90% (± 8.51%) of the reads mapped to the *B. rapa* genome on average. In total, 2.05 million SNPs and 0.11 million indels were identified, with 13,760 multi-allelic SNP sites, average SNP depth 4.69 (± 1.97), and a transition/transversion ratio of 1.34. After filtering, 31,662 high quality SNPs were retained for our phylogenetic and genetic structure analyses and 426,740 SNPs were retained for our demographic analyses. A summary of SNPs on each chromosome is available in Table S3.

### Inference of Genetic Structure and Phylogenetic Relationships

Our genetic structure and phylogenetic analyses of 31,662 SNPs supported five distinct genetic clusters among the 126 *B. rapa* accessions. Analyses of genetic structure recovered six clusters using fastStructure (Fig. 1, Fig. S1). Increasing or decreasing the number of clusters by one did not substantially change the observed clustering pattern (Fig. 1, Fig. S1), and *K=* 6 clusters maximized the marginal likelihood (Fig. S2). Most of the morphologically distinct crops long recognized as subspecies—pak choi, Chinese cabbage, rapini, and the yellow sarson—were largely resolved as distinct clusters in our fastStructure analyses. Notably, they are also well resolved with *K* = 5 or 7 clusters (Fig. 1). We also identified a number of genetically diverse accessions (multi-color in fastStructure results, Fig. 1). Some of these accessions are recently admixed crop lines from the US, but the history of many of these accessions is unclear. The sixth cluster (gray color on Fig. 1) resolved in the fastStructure *K* = 6 analysis described genetic variation in these diverse accessions. Otherwise, fastStructure resolved five largely homogenous clusters (Fig. 1). Analyses with CLUMPAK found that the same genotypes were consistently assigned to these clusters across ten iterations of fastStructure (Fig. S3).

The phylogenetic relationships of the *B. rapa* accessions were well resolved and consistent with our fastStructure clustering. More than 98% of the nodes on the RAxML tree had bootstrap support higher than 50% (Fig. S2). Many of the morphologically distinct crops were distinguished as well supported clades in our phylogenomic analysis. The combined phylogenetic and genetic clustering analyses resolved five major genetic groups within *B. rapa* that largely corresponded to different crops. These groups were a European-Central Asian cluster (group “EU-CA”) represented by *B. rapa, a* group that contained rapini and other accessions (group “S”) represented by *B. rapa* subsp. *sylvestris* (bootstrap support 100%), an Indian sarson group (group Ί”) comprised of *B. rapa* subsp. *trilocularis* and three accessions of subsp. *dichotoma* (bootstrap support of 100%), a chinensis group (group “C”) composed of *B. rapa* subsp. *chinensis* (bootstrap support of 68%), and most of the *B. rapa* subsp. *pekinensis* accessions clustered together in a *pekinensis* group (group “P”; bootstrap support of 66%). Interestingly, the eight *B. rapa* subsp. *oleifera* (blue stars on Fig. 1 and Fig. S2), six *B. rapa* subsp. *rapa* (pink stars on Fig. 1 and Fig. S2) and five of the *dichotoma* accessions (green stars on Fig. 1 and Fig. S2) were distributed across the phylogeny. Genetically heterogeneous accessions identified in our fastStructure analyses were scattered across the phylogeny. Groups of these heterogenous samples were sister to each of the five largely homogenous genetic groups. Whether these heterogeneous accessions are the result of recent admixture or represent ancestral variation is not clear from the current analyses.

The phylogenetic analyses also supported a European-Central Asian origin for *B. rapa.* Using two *B. oleracea* accessions as an outgroup, the concatenated phylogenetic analysis of 31,662 SNPs resolved the European-Central Asian *B. rapa* cluster (EU-CA) as sister to all other *B. rapa* accessions (Fig. 1; Fig. S2). Included in this clade were most of the European and Central Asian accessions we sampled *(B. rapa).* This relationship was well supported with a bootstrap value of 71%. The nested set of sister relationships among the remaining taxa indicated an eastward expansion of *B. rapa* crops. Accessions from East and South Asia were among the most derived lineages in our phylogeny.

### Demographic History of *Brassica rape’s* Domestication Across Asia

Our ∂ a ∂ i analyses of the demographic history of *B. rapa* also supported an eastward series of domestication events over the past few thousand years (Fig. 3 and Table 1). The demographic models were well supported qualitatively: the chosen optimal project size maximized the number of alleles captured in SFS under each model (Fig. S4); and the best fit model for each group reproduced the observed frequency spectra with no severe deviations in the residuals (Fig. 4). Across the range of *B. rapa,* our data supported a stepwise eastward progression of crop domestication over the past 2446-4070 years (Fig. 3b and Table 1) and were best explained by a model that involved an initial population diverging from EU-CA, followed by SA split with EA (Fig. 2a, SA v EA 3; Fig. 3a). This model was supported with a much higher log-likelihood than the next best fitting models (Δ =2469; log-likelihood = -16390). Under this model, we inferred the time span from initial divergence of SA-EA to the split of SA-EA was approximately 976-1630 years, and the divergence of EA and SA occurred approximately 1470-2440 years ago (Table 1, Fig. 3).

**Fig. 3.**
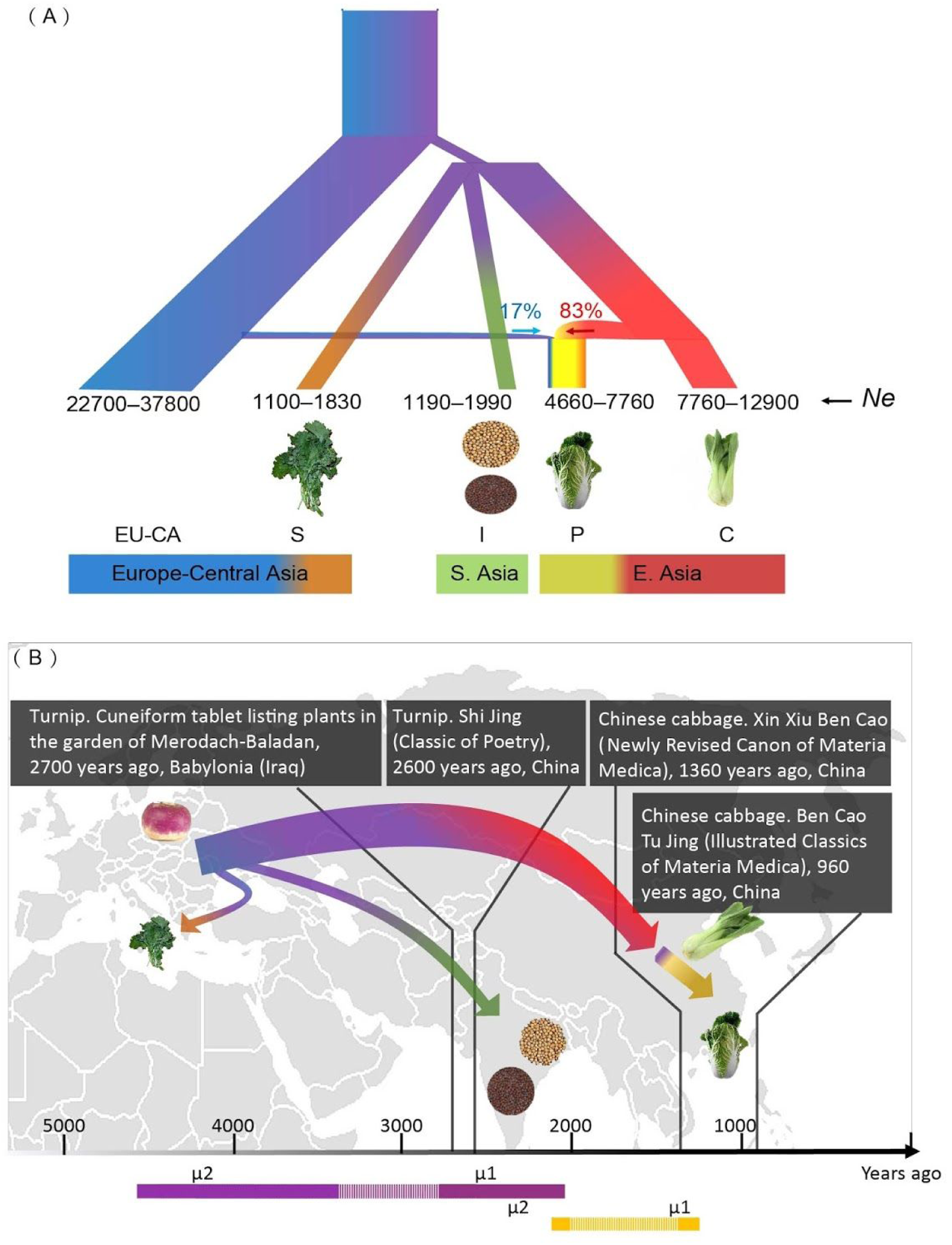
A combined summary of demographic inferences and written records of *B. rapa* domestication events. **(a)** The combined best-fitting demographic models based on ∂ a ∂ i simulations with absolute parameter values. The colored parallelograms represent different *B. rapa* genetic groups: European-Central Asian *B. rapa* and *B. rapa* subsp. *rapa* (purple); *B. rapa* subsp. *sylvestris* (orange); *B. rapa* subsp. *trilocularis* and *B. rapa* subsp. *dichotoma* (green); *B. rapa* subsp. *pekinensis* (yellow); *B. rapa* subsp. *chinensis* (red). The width of the branches represents relative *Ne*. The numbers on the parallelograms represent the effective population size estimates (unit: individual). A horizontal band represents an admixture event with proportions defined by parameters in the model, **(b)** A plot of the eastward introduction and diversification of *B. rapa* with a time scale of key domestication events corroborated by historical written records. The horizontal arrow represents time scale. Gray callouts identify the historical written records of *Brassica rapa* crops (cultivar, source, time, country). The purple and yellow solid bars represent the estimated time of eastward introduction and domestication of Chinese Cabbage in ∂ a ∂ i based on two substitution rates (μ1 and μ2), respectively. The dashed bars represent the time range between the inferences from two different substitution rates.

**Fig. 4.**
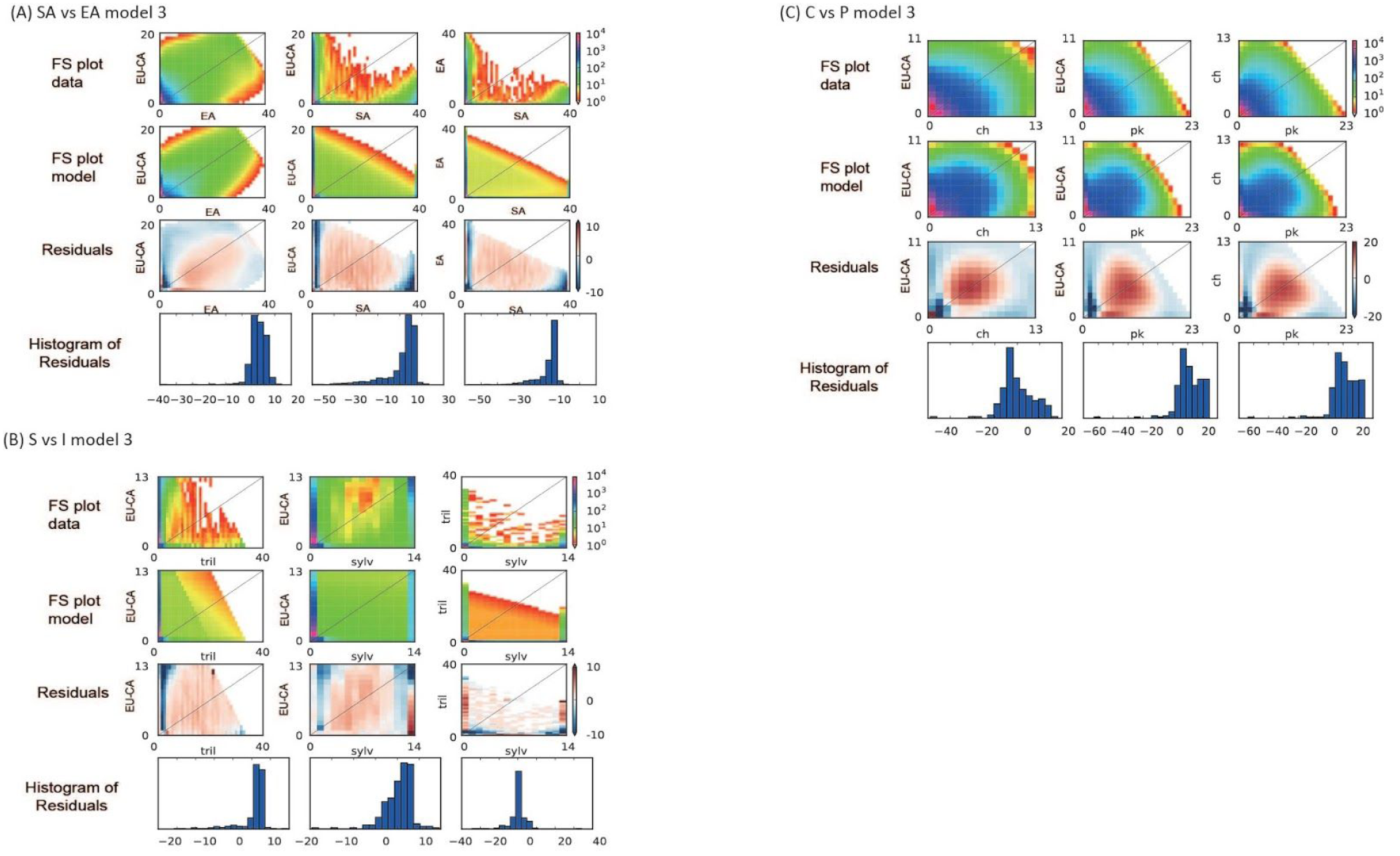
Population pairwise plots of 3D model-data comparison for each optimal model as logarithmic color-maps under the tested models. (A) EA vs SA 3; (B) S vs I 3; (C) C vs P 3. For each model, the first row is the data frequency spectrum plots, the second row is the model frequency spectrum plots. On each heat plot, the upper-left is the model and data frequency spectra, the lower-right is the residuals between model and data. The third and fourth row are the plots of residuals and histograms of residuals. Dark red or blue on the residual plots indicate the model predicts too many or few alleles. The residual patterns of ∂ a ∂ i modeling could be explained by ongoing gene flow between the populations, which was not included in our demographic models.

Among the models exploring the relationships of the primarily European, Central Asian, and South Asian accessions (Fig. 2b), our analyses supported an initial common ancestor split from EU-CA followed by subsequent divergence into the *sylvestris* (“S”) and *trilocularis + dichotoma* (Ύ) groups (S v I 3). The log-likelihood value of this model (-4959) is much higher than that of the other two alternative models (Δ =12280). Under this optimal model, a population split from EU-CA approximately 3715-6190 years ago (Table 1). This population subsequently split into groups S and I (T1 under S v I 3 in Table 1 and Fig. 2) 825-1380 years later. This time span before divergence of later groups is consistent with the corresponding estimate for the duration of a common ancestral population in the optimal model of model group A (SA vs EA 3; 976-1630 years). Under this model, we inferred the split of S and I occurred 2890-4810 years ago.

Our simulations on the origins of Chinese cabbage (*B. rapa* subsp. *pekinensis)* supported an ancient admixed origin of the crop. The optimal demographic model under indicated that pekinensis was formed by admixture of *chinensis* and *rapa* (model C v P 3, Fig. 2c). This optimal model had a log-likelihood value (-18110) much higher than that in the two alternative models (Δ = 383). An estimated 83% of the *pekinensis* genetic variation came from *chinensis,* whereas the remaining 17% was derived from European-Central Asian *rapa.* Admixture that contributed to the founding of *pekinensis* is estimated to be ancient. The best fitting demographic model (Table 1) estimated the admixture occurred 1244-2070 (± 52-86) years ago.

Our hierarchical model testing in ∂ a ∂ i also allowed us to compare the relative effective population sizes (Λ/e) of the *B. rapa* genetic groups. Given *Ne*_EU-CA_ was the reference standard for the derived populations under each model, and the estimated *Ne* values for each EU-CA group were similar across the three model groups (*Ne*_EU-CA_ = 22700-25900 if μ_1_ = 1.5 × 10^−8^ or *Ne*_EU-CA_ = 37800-43100 if μ_2_ = 9 × 10^−9^; Table 1), it is reasonable to compare the estimated *Ne* for each group across models. Under the “S v I 3” and “C v P 3” ∂ a ∂ i analyses, we observed four to seven times larger *Ne* in *chinensis* (*Ne_c_* = 7760-12900) and *pekinensis* (*Ne*_P_ = 4660-7760) than that in *sylvestris* (*Ne*_S_ = 1100-1830) and *trilocularis* and *dichotoma* (Ne_l_ = 1190-1990). This pattern is consistent with the results based on model “EΑ v SA 3” under which *Ne*_EA_ is approximately 13 times larger than *Ne*_SA_.

### Nucleotide Diversity of the *B. rapa* Genetic Groups

Finally, we also compared the genetic diversity of the five genetic groups of *B. rapa* identified in our analyses. The mean nucleotide diversity of the *sylvestris* group (Π_S_ = 1.08 × 10^−5^) and Indian *trilocularis* and *dichotoma* sarson group (π, = 0.97 × 10^−5^) were 63% and 66% lower than EU-CA group (Π_EU-CA_ = 2.88 × 10^−5^). In contrast, the *chinensis* (Π_C_ = 2.45 × 10^−5^) and *pekinensis* (Π_ρ_ = 2.40 × 10^−5^) groups did not have a similarly strong decrease of nucleotide diversity (15% and 17% lower) relative to the EU-CA group (Fig. 5). Together, our ∂ a ∂ i analyses and nucleotide diversity analyses suggested a less severe genetic bottleneck with at least one ancient admixture event in the East Asian group relative to the South Asian group during the eastward progression of *B. rapa.*

**Fig. 5.**
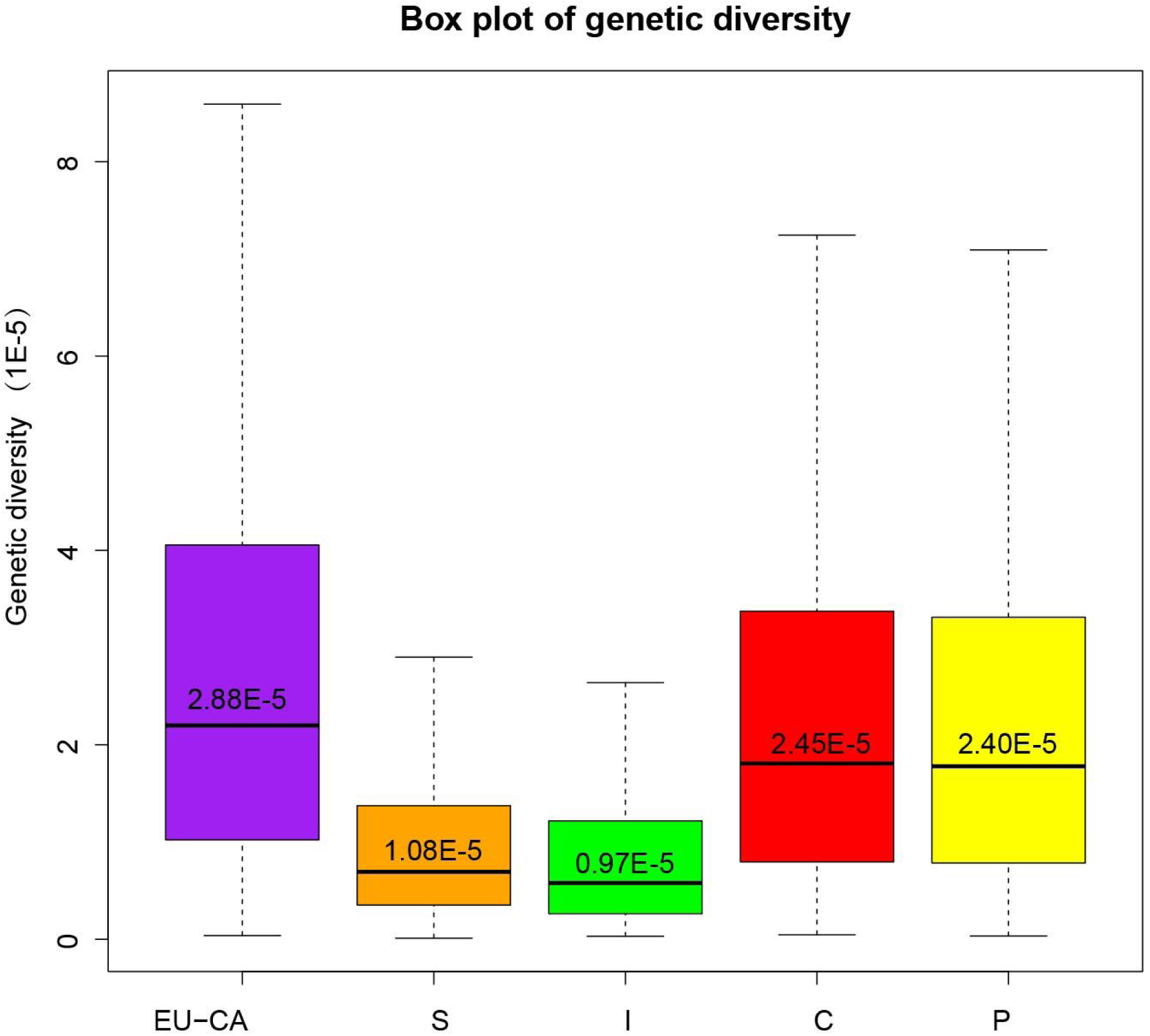
Box plot of nucleotide diversity of the five *B. rapa* genetic groups. Bottom and top of the box represent the first and third quartiles. Band inside the box represent median. The numbers represent mean values for each group. Colors of each group are consistent with fastStructure results. Abbreviation: EU-CA represents *B. rapa* and *B. rapa* subsp. *rapa* group; S represents *B. rapa* subsp. *sylvestris* group; I represents *B. rapa* subsp. *trilocularis* and *B. rapa* subsp. *dichotoma* group; C represents *B. rapa* subsp. *chinensis* group; P represents *B. rapa* subsp. *pekinensis* group.

## Discussion

Our analyses of more than 2 million SNPs across 126 newly sequenced genotypes of *Brassica rapa* clarified the history and origins of multiple *Brassica* crops. We found strong evidence for at least five genetic groups within *B. rapa* from our combined phylogenetic, genetic structure, and demographic analyses. Most previous studies identified two or three geographically based groups: Europe-Central Asia and East Asia (Zhao *et al.* 2005; Takuno *et al.* 2006), and South Asia if sampled (Song *et al.* 1990; Del Carpio *et al.* 2011a; b; Guo *et al.* 2014). Our results are largely consistent with these analyses. However, we found more resolution of genetic structure among East Asian crops and recognized a genetically distinct cluster that contains *B. rapa* subsp. *sylvestris.* These results are also consistent with a recent genomic analysis that used ˜20,000 SNPs to study *B. rapa* relationships (Cheng et al. 2016). Although our sampling differs from Cheng et al. (2016) —we sampled rapini (subsp. *sylvestris*) and brown sarson (subsp. *dichotoma*), more yellow sarson (subsp. *trilocularis*), and more diverse European and Central Asian accessions—both of our analyses recovered Chinese cabbage, pak choi, and a European-Central Asian group that includes European turnips. Their sampling included more Asian crops than our analyses, and they resolved additional groups that included caixin. (subsp. *parachinenesis*), zicatai (subsp. *chinensis var. purpurea*), and Japanese turnips with mizuna (subsp. *nipposinica*). In contrast, we sampled many accessions not previously analyzed with genomic data, including rapini (subsp. *sylvestris*) and the Indian sarsons (subsp. *dichotoma* and *trilocularis*), and recovered distinct genetic groups associated with these accessions. Further, our more sophisticated modeling of the demographic history of *B. rapa*—corroborated by written records—provides new insights on the age and relationships among the diverse *B. rapa* lineages. Considering the age estimates of population splits among the *B. rapa* groups, it is not surprising that past analyses with fewer markers were unable to resolve the genetic structure among these plants. Although our analyses provide strong evidence for at least five groups, not all individuals of *B. rapa* could be placed in one of these distinct genetic clusters. Determining whether this is due to parallel evolution, admixture (recent or ancient), insufficient sampling of related genotypes, or lack of divergence requires further research with new samples. Regardless, our analyses, as well as those of others (Song *et al.* 1990; Zhao *et al.* 2005; Takuno *et al.* 2006; Del Carpio *et al.* 2011a; b; Guo *et al.* 2014; Cheng *et al.* 2016), contrast with one recent study (Tanhuanpää *et al.* 2016) and find evidence that geography and crop type contribute to the genetic structure of *B. rapa.*

One of the interesting aspects of the domestication of *B. rapa* crops is that many were domesticated during written history. This provided an opportunity to compare our population genomic results to historical records for many of the demographic splits modeled with ∂ a ∂ i. Previous simulations of ∂ a ∂ i’s accuracy found that inferences of divergence times had low error with even modest sample sizes (Robinson *et al.* 2014). A recent empirical analysis using ∂ a ∂ i recovered evidence of known recent demographic changes in the checkerspot butterfly (McCoy *et al.* 2014). Our inferred times of demographic events in the history of *B. rapa* are consistent with these previous analyses and appear to be accurate as they were consistent with the available written record. Demographic inferences based on genome wide SNP data found that the eastward introduction and diversification of *B. rapa* happened 2400-4100 years ago. This inferred time span is compatible with the two oldest historical records of *B. rapa* in East and South Asia. *Brassica (Li 1981; Ye 1989; Luo 1992)* was first described in the ancient Aryan literature of India nearly 3500 years ago (Prakash *et al.* 2011), and in Shi Jing (Classic of Poetry), the oldest existing Chinese classical poetry collection (Li 1981; Ye 1989; Luo 1992). Similarly, our ∂ a ∂ i analyses estimated Chinese cabbage (*B. rapa* subsp. *pekinensis*) was developed 1200-2100 years ago. The earliest historical record of Chinese cabbage is nearly 1400 years old. Niu Du Song (i.e., beef tripe cabbage) is mentioned in Xin Xiu Ben Cao (Newly Revised Canon of Materia Medica), one of the oldest official pharmacy books edited by Jing Su in 659 CE. A clear description of Chinese cabbage is also found in Song Su’s Ben Cao Tu Jing (Illustrated Classics of Materia Medica) from 1061 CE (Li 1981). He described Niu Du Song in Yangzhou city in Southern China as a plant with large, round, crinkled leaves and a less fibrous texture. Beyond the timing of these events, the inferred order of domestication is also supported by the written records. Our results suggests that turnips are likely the earliest domesticated crop with leaf and seed crops domesticated later. A translated passage in the Shi Jing supports this order: when collecting turnip and radish, we should not abandon them them because of their bitter root because the leaves and stems taste good. These written records corroborate our population genomic inference of the timing of two different events in the history of *B. rapa* domestication and lend confidence to the other estimated parameters.

Notably, the inferred timing of these demographic events better overlapped with estimates from one of the Brassicaceae substitution rates. substitution rates in the Brassicaceae have recently been debated as a new fossil provided a different calibration for the family (Beilstein *et al.* 2010). The placement of this fossil has been the subject of debate (Franzke *et al.* 2016) because phylogenetic inferences generally yield older dates for macroevolutionary events (Beilstein *et al.* 2010; Arias *et al.* 2014) than previous estimates (Koch *et al.* 2000; Barker *et al.* 2009; Edger *et al.* 2015; Hohmann *et al.* 2015). To work within this debate, our demographic estimates were calculated twice using two different estimates of the Brassicaceae nuclear substitution rate: the faster rate from Kock et al. (2000) and a more recent, slower rate developed by Kagale et al. (2014) based on the fossil placement of Beilstein et al. (2010). Three of the four written records occurred within the 95% confidence interval for the timing of population splits estimated with μ_1_ whereas all of the estimates using μ_2_ preceded the available written records. Thus, both of these substitution rates are consistent with the written record although there is greater concordance with the more recently derived and slightly slower substitution rate. Considering that we expect a written record to follow a domestication event with some lag, the “true” substitution rate for *Brassica* and other Brassicaceae may lay somewhere between these two rate estimates.

The ∂ a ∂ i modeling also indicated that the South Asian group (“S” and “I”) experienced a more severe founder effect than the East Asian group (“EA”). The estimated effective population size and nucleotide diversity of the East Asian group were several-fold larger than the South Asian group. This difference of founder effect within a single species is striking in comparison to other crops. The South Asian group retains only approximately 35% of the genetic diversity of the EU-CA group. This is equivalent to the domestication bottleneck in barley (Caldwell *et al.* 2006), rice (Zhu *et al.* 2007; Caicedo *et al.* 2007), and among the most severe bottlenecks found in annual crops (Miller & Gross 2011). In contrast, the East Asian group retained approximately 84% of the genetic diversity present in the EU-CA group. The domestication bottleneck of the East Asian *B. rapa* crops is comparable to the bottleneck in sorghum (Casa *et al.* 2005), sunflower (Tang & Knapp 2003), soybean (Li *et al.* 2010), and among the least severe bottlenecks found in annual crops (Miller & Gross 2011). Considering the two lineages diverged only a few thousand years ago and both likely experienced long-distance introductions from ancestral European-Central Asian populations, the difference of *Ne* and π between the two groups is probably due to a combination of differences in gene flow (Meyer & Purugganan 2013) and inbreeding (Dempewolf *et al.* 2012). In East Asia, commercial activity along the ancient silk-road (started 220 BC) may have continuously brought diverse *B. rapa* genetic resources to East Asia. Consistent with this hypothesis, we found evidence of shared genetic variation among some European-Central Asian and East Asian accessions, as well as a small amount of EU-CA admixture in the origin of *B. rapa* subsp. *pekinensis.* In contrast, South Asian *B. rapa* may have been more punctually introduced with events such as Alexander the Great’s invasion of India in 325 BCE. Written records document his troops likely brought mustard seeds as army provisions (Grubben 2004). A shift to predominant inbreeding in the yellow and brown sarsons (McGrath & Quiros 1992) likely further reduced their genetic diversity. Thus, domestication of these two different regional groups of *B. rapa* crops occurred with significantly different population genetic backgrounds that essentially spans the range of possible models of domestication (Eyre-Walker *et al.* 1998; Miller & Gross 2011; Meyer & Purugganan 2013; Gaut *et al.* 2015). This less severe founder effect in the EA group may contribute to the diverse crop types in East Asia, in contrast to the Indian group.

Despite the economic importance of *B. rapa,* the order of crop domestication has never been well resolved. Previous studies generally considered *B. rapa* subsp. *sylvestris* to be the wild type and reflect a European *B. rapa* center of origin (Guo *et al.* 2014; Tanhuanpää *et al.* 2016). However, *B. rapa* subsp. *sylvestris* has never been genetically distinguished from other *B. rapa* subspecies. Genome-wide analyses with 715 polymorphic SSR alleles (Guo *et al.* 2014) and 209 nuclear SNPs (Tanhuanpää *et al.* 2016) grouped *B. rapa* subsp. *sylvestris* among European accessions or clustered with oil types. A recent whole genome resequencing study of *B. rapa* (Cheng et al 2016) did not include any *sylvestris* accessions. Our results suggest *sylvestris* is not likely a wild *B. rapa.* The two *sylvestris* accessions and its five genetically cognate accessions in our analyses clustered into a monophyletic clade that was supported with 100% bootstrap support in our phylogenetic analysis. These accessions were also grouped in our genetic structure analyses and present in multiple values of *K.* Importantly, this group was not sister to all of the remaining *B. rapa* accessions. Using two *B. oleracea* as outgroups to root the phylogenetic tree of *B. rapa,* we found a collection of European and Central Asian *B. rapa* accessions were sister to the remaining *B. rapa.* Based on recent whole genome resequencing analyses (Cheng *et al.* 2016), previous linguistic evidence (Ignatov *et al.* 2008), and archaeological records (Körber-Grohne 1987), turnip is likely the first domesticated *Brassica* in the European-Central Asian region. Our study is less clear on this question as only one of the turnip accessions was placed in the EU-CA clade. However, none of our turnip collections are from the European-Central Asian region, and 2/3 of them have a complex genetic background based on our fastStructure analyses. Our result is congruent with recent reports by Cheng et al. (2016) that there may be two turnip lineages (European and Asian turnip). Further sampling of turnips from across the geographic range would address many of the remaining questions surrounding the early domestication of *B. rapa.*

With an expanded sample of yellow sarson and brown sarson, we also found evidence for a single origin of yellow sarson (*B. rapa* subsp. *trilocularis*), but possibly multiple origins of brown sarson (*B. rapa* subsp. *dichotoma*). All accessions of *trilocularis* and three of the *dichotoma* accessions formed a monophyletic clade in our phylogenetic analysis. This is consistent with a single origin of the yellow sarsons with multiple origins of brown sarsons from yellow sarsons. Further, a couple individuals of *dichotoma* clustered more closely with *sylvestris* and a single individual was grouped with *chinensis.* Other *dichotoma* individuals were distributed among the genetically heterogeneous accessions. Assuming that our accessions are correctly labeled, these results suggest that brown sarsons may have more than one origin from non-sarson ancestors. It is worth noting that the USDA nomenclature system groups brown sarson together with another crop, toria, as subsp. *dichotoma.* This may be the cause of apparent polyphyly of subsp. *dichotoma.* Alternatively, the placement of some *dichotoma* accessions may reflect more recent admixture rather than independent origins. These results are mostly consistent with a previous analysis based on five isozyme and four RFLP markers (1992). Other similar *B. rapa* population and phylogenetic studies typically adopted or mentioned these two subspecies, but only included a few samples to draw phylogenetic inferences (e.g., Cheng *et al.* 2016 and Del Carpio *et al.* 2011b). Thus, our results indicate that the potential diversity and relationships of these morphologically similar crops have yet to be fully explored. Additional sampling beyond the available USDA accessions of the yellow and brown sarsons—including both self-compatible and incompatible types as well as clearly identified representatives of toria—is needed to further resolve the origins and diversity of these crops.

Oil crops are among the most economically important crops of *B. rapa,* and we found evidence for multiple origins of oil type *B. rapa* subsp. *oleifera.* The eight *oleifera* samples were distributed throughout our phylogeny and occurred in four of the five major genetic groups. Considering that by definition canola is a *Brassica* species containing less than 2% erucic acid and less than 30 micromoles glucosinolates (Eskin & McDonald 1991), it is not surprising to find *oleifera* samples scattered across the phylogeny. Oil crops are recognized as being genetically and geographically diverse relative to other cultivated crops because of likely multiple origins of oilseed crops (McGrath & Quiros 1992; Takuno *et al.* 2006; Tanhuanpää *et al.* 2016). Our results also support parallel origins of the oil types of *B. rapa* from different genetic backgrounds. Oil type *B. rapa* appears to have been primarily selected from mostly non-leafy crops with a concentration among the European-Central Asian genetic group.The oil crops of *Brassica* may have multiple, independent origins. Studies on oilseed rape (*Brassica napus*) have also shown that they have relatively diverse accessions that have been domesticated for winter and spring oilseed rape in Europe and Asia(Allender & King 2010; Li *et al.* 2014; Wu *et al.* 2014; Gazave *et al.* 2016). Considering that most *Brassica* oil cultivars have been domesticated relatively recent (Reiner *et al.* 1995) that post-date the globalization of agriculture (Phillips & Khachatourians 2001; Font *et al.* 2003; Kumar *et al.* 2015), it is not surprising that they were sampled from throughout the phylogeny. Thus, the domestication of oil type *B. rapa* may similar to the parallel domestication of those oil types in *B.napus.*

Our relatively large sampling and demographic analyses allowed us to resolve a long-held debate on the origin of Chinese cabbage. Previous genetic and empirical studies supported two distinct hypotheses for the origin of Chinese cabbage: directly from pak choi (Song *et al.* 1990; Zhao *et al.* 2005; Takuno *et al.* 2006) or via genetic admixture of turnip and pak choi (Li 1981; Ren *et al.* 1995). Our fastStructure and phylogenetic inference suggest *pekinensis* has a closer relationship with *chinensis.* Our demographic modeling supported an admixed origin of *pekinensis* via crossing of EU-CA *Brassica rapa* and subsp. *chinensis,* with *chinensis* contributing ~90% of the contemporary *pekinensis* genome. Given that most of the genome is *chinensis,* it is not unexpected that previous analyses with fewer markers would not detect the modest 10% genomic contribution from European-Central Asian *B. rapa.* The fastStructure results may also indicate the *B. rapa* subsp. *rapa* accessions most closely related to the progenitor of *pekinensis.* Components of genetic variation in a handful of *B. rapa* subsp. *rapa* accessions were assigned to the same category as the bulk of our *pekinensis* genotypes. It is not clear from the fastStructure analysis if this structure represents the ancestral variation contributed by *rapa* to *pekinensis,* or more recent admixture from *pekinensis* into some *rapa* accessions. Detailed analyses and expanded sampling are required to further dissect the contributions and history of *B. rapa* subsp. *pekinensis.* Our analyses also demonstrate that Chinese cabbage was formed much later than other Asian *B. rapa* crops and indicate it is among the most recently derived *B. rapa* subspecies. Samples of other East Asian leafy type subspecies (*parachinensis, narinosa, perviridis* and *nipposinica*) were placed in either the *pekinensis* or *chinensis* clades, consistent with a recent study with abundant sampling of these leafy types (Cheng *et al.* 2016).

## Conclusions

Overall, our study provides many new data and insights into the history of *B. rapa* and provides clear direction for future research. Our expanded sampling of frequently underrepresented groups, such as the Indian sarsons, has provided evidence on their origins and diversity. Expanded sampling of genetic diversity within *B. rapa* will likely resolve additional groups and clarify the complex history of these plants. In particular, better genomic sampling of *B. rapa* landraces from Central Asia, landrace turnip from Europe, landraces and elite rapini from Italy, landrace toria, and oil type *oleifera* are needed to further resolve the domestication history of *B. rapa.* These samples would augment the existing collections of *B. rapa* in North America and aid curation of the existing USDA collection which contained many misidentified accessions. Sequence data from wild *B. rapa* seeds or archeological materials would also significantly improve our analyses. Additional genomes from putative wild *B. rapa* populations would also mitigate ascertainment bias in future analyses and provide more insight into changes in structural variation associated with domestication. Despite potential for further improvement, our results have provided the most robust evidence to date on the genetic structure and domestication history of *B. rapa.* Our use of demographic modeling provided some of the first population genomic estimates of the ages and population size changes that occurred during *B. rapa’*s history. Corroboration of the estimated dates with the written record provides added confidence in our inferences. Given that the other *Brassica* crops are of similar age with equally complicated histories, the population genomic approach employed here will likely provide new insights into *Brassica* domestication when applied to the other crops. Most importantly, our results provide a new framework for *B. rapa* that includes empirical estimates of domestication times, effective population sizes, and the amount and direction of gene flow between divergent lineages.

## Acknowledgements

We thank A. Baniaga, K. Dlugosch, S. Jorgensen, H. Marx, and L. Zheng for helpful comments on the manuscript. Server hosting infrastructure and services provided by the Bio Computing Facility (BCF) at the University of Arizona. X.Q., T.E.H., H.A., J.C.P., and M.S.B. were supported by NSF-IOS-1339156. A.R. and R.G. were supported by NSF-DEB-1146074.

## Data Accessibility

SRA files for the samples are available at NCBI-SRA database (SRP072186, http://www.ncbi.nlm.nih.gov/sra/SRP072186).

## Author Contributions

M.S.B., J.C.P. and R.G. designed the experiments. X.Q., H.A., A.R. and T.E.H performed the experiments, X.Q. and M.S.B. wrote the paper.

## Supplementary Tables and Figures

Table S1 Sample information of the investigated *Brassica rapa* accessions.

Table S2 A list of the misidentified *Brassica* seeds from USDA. None of these accessions were used in this study.

Table S3 Summary of SNPs on each chromosome using *Brassica rapa* genomes version 1.5 as reference.

Table S4 Image sources.

Fig. S1 Marginal likelihood distribution of the fastStructure clustering with *K* ranging from 1 to 10. Python script *chooseK* coming with the fastStructure package was used to estimate the best *K* value. *K* = 6 explained the population structure best and maximized marginal likelihood.

Fig. S2 The integrated maximum likelihood phylogeny and fastStructure results of the 126 investigated *Brassica rapa* accessions. An expanded version of Fig 1 with detail accession information.

Fig. S3 A comparison of fastStructure result and CLUMPAK result based on ten iterations when *K=* 6.

Fig S4 The trend lines illustrate the relationship between project size and the number of alleles captured in SFS under model (a) and model (b, c).

